# Surface-by-temperature interactions shape the relationship between viral free-living survival and reproduction

**DOI:** 10.1101/2025.11.16.688743

**Authors:** Ketty Kabengele, Anmol Seth, Thomas Johnson, C. Brandon Ogbunugafor

## Abstract

Environmental factors play a critical role in viral persistence, yet the combined effects of temperature and surface chemistry on viral fitness are not fully understood. We examined bacteriophages T4 and *φ*X174 across copper, stainless steel, and polypropylene plastic surfaces at 4°C and 37°C to assess how these variables influence survival and reproduction. Both phages exhibited marked differences in survival across surfaces: copper at 37°C caused rapid inactivation, and cooler conditions promoted persistence. Reproductive capacity varied independently of survival, indicating that viral stability and replication are partially decoupled traits shaped by environmental context. Linear modeling and correlation analyses confirmed significant, non-additive interactions between surface type and temperature. These findings suggest that abiotic factors can act synergistically to determine viral fitness outside the host. More broadly, they refine our understanding of the mechanisms that govern the environmental ecology of viruses and the canonical relationship between survival and transmission.

## 1 Introduction

The quest to understand how pathogens persist and transmit is a central goal of infectious disease ecology. Pathogens employ diverse strategies to move between hosts, including airborne, asymptomatic, sexual, vector-borne, and others (*1,2*). Among the least studied of these pathways is indirect transmission (*3*), where pathogens are transmitted through an environmental intermediate, such as a physical surface, sometimes summarized as transmission via “fomites” (*4–11*).

In viruses, free-living survival on surfaces has been the focus of growing research interest, as persistence in the environment can influence opportunities for infection. For example, studies of influenza (*12–17*), noroviruses (*18–27*), and coronaviruses (*28–33*) have demonstrated that viruses differ in their ability to survive on surfaces, which may influence outbreak dynamics and public health interventions. Metals such as copper are known to exhibit strong antimicrobial activity (*34–41*), while materials such as stainless steel and plastics often preserve viral integrity. In addition to the study of surface survival, previous work has identified several abiotic factors that influence viral survival, including temperature, humidity, and pH (*30,36,42–45*). Similarly, cooler temperatures tend to extend viral persistence relative to warmer ones (*43,46,47*). A larger literature on thermal adaptation in viruses has revealed that temperature adaptation can be a highly evolvable trait across different virus types (*48–50*). Together, these findings highlight the importance of abiotic stressors in shaping viral persistence. However, most studies have examined abiotic factors in isolation. Less well understood are the interactive effects of multiple stressors operating in tandem.

An additional dimension often overlooked in studies of viral persistence is reproduction. Survival on surfaces is typically treated as synonymous with fitness, and scientists have often framed these traits in the framework of life history theory (*51–56*). Furthermore, experimental and theoretical studies have proposed that survival and reproductive potential can have a variety of relationships, depending on the characteristics of virus-host interactions (*55,57–59*). This raises fundamental questions about how survival and reproduction relate in different environments and whether the relationship between these traits is consistent, context-dependent, or idiosyncratic.

Modern epidemiology underscores the importance of these questions. During the COVID-19 pandemic, concerns about the surface transmission of SARS-CoV-2 via fomites led to global efforts to disinfect surfaces, and studies showed that the virus could persist for hours to days depending on temperature, humidity, and surface material (*28,60,61*). Although later work confirmed that airborne transmission was the dominant route, surface stability remains a key consideration for coronaviruses and other respiratory pathogens. Studies of environmental persistence and free-living survival were also important during recent outbreaks of the Mpox virus (MPXV). Studies have demonstrated that MPXV exhibits remarkable stability on environmental surfaces, with viable virus detected on household items at least 15 days after contamination (*62*). In addition, MPXV demonstrates temperature-dependent persistence, with longer survival at lower temperatures (*63*). Importantly, the type of surface significantly influences the persistence of MPXV, as porous materials such as bedding and clothing harbor viable virus more than non-porous surfaces (*62*). The stability of MPXV even varies substantially depending on the biological matrix, with the virus showing increased persistence in proteinaceous fluids such as blood and semen (*64*). These findings collectively highlight how understanding specific environmental conditions influences both survival and reproductive potential, knowledge that can help refine public health strategies, improve risk assessment and inform theoretical frameworks that guide constraints on pathogen evolution.

To address outstanding questions about how environmental factors affect the relationship between viral free-living survival and transmission, we investigated the manner in which temperature and surface composition jointly influence viral survival and reproduction. Using two well-characterized bacteriophages, *φ*X174 (*65*) and T4 (*66*), as model systems, we tested survival and reproductive efficiency on three surfaces (copper, polypropylene plastic and stainless steel) at two temperatures (4°C and 37°C). These phages provide useful models for general principles of viral ecology because they are genetically distinct, differ in morphology and life history strategies, and have been widely used in laboratory-based studies of stability and evolution.

By quantifying decline rates, reproductive efficiency, and correlations between survival and reproduction across surface-temperature environments, our study provides insight into how abiotic interactions shape viral fitness. More broadly, this work demonstrates how combinations of environmental contexts can drive non-additive outcomes in viral ecology, with implications for understanding the persistence of human and animal pathogens, informing epidemiological models of fomite-mediated transmission, and informing theories of microbial ecology and epidemiology.

## 2 Materials and Methods

### 2.1 Bacteriophage propagation and host growth

Both *φ*X174 (ATCC 13706-B1) and T4 (11303-B4) phages were purchased from the American Type Culture Collection (ATCC) and grown in their recommended host bacteria, *E*.*coli. φ*X174 was grown in the host bacteria *E*.*coli* strain C (ATCC BAA-13706) following the ATCC guidelines, and T4 was grown in MG1665 derived from *E*.*coli* K-12 (*67*).

Both hosts were grown overnight at 37°C with shaking at 160 rpm in their respective media (MG1655 in Luria-Bertani liquid media, *φ*X174 host in nutrient broth (ATCC Medium 129). Viral propagation was performed in liquid media according to protocols recommended and published on the ATCC website. Viral dilutions were performed in appropriate liquid media, and different concentrations were tested to determine the most suitable concentrations: 10^−5^ for *φ*X174 and 10^−7^ for T4. All dilutions were stored at 4°C as recommended by ATCC. All material surfaces were purchased online from Amazon. High-density polypropylene plastic (USA Sealing bulk-PS-PP-323, 1/8 inch in height, 16 inches in width, 16 inches in length) was purchased and cut into smaller squares (2 inches long and 2 inches wide). Stainless steel discs (2 inches in diameter, 15 gauge thickness circular plate 304), and circular copper metal discs (24.26 mm in diameter, 99.9 % pure). All materials were thoroughly sanitized, washed, and soaked in alcohol prior to the experiments.

The two viruses were exposed to these surfaces at 4°C and 37°C. The controls were performed in Eppendorf tubes at 4°C. 2000 µL of phage lysate was added to the appropriate surface for eight hours, and samples were taken every two hours for viral quantification via plaque assay (**Figure** 1(a)). At the final time point (8 hours), samples were taken from each surface, and reproduction was measured by infecting bacteria with the sampled phages (**Figure** 1(b)). Reproduction was measured as the number of phages reproducing within an hour, starting at time 0, and was measured every 10 minutes for 30 to 60 minutes. Viral quantification was performed using the plaque assay method. 100 µL of appropriate host bacteria were mixed with 100 µL of phages, and 3 mL of 50°C top agar (25g of LB broth, 7.5g agar, and 1L of DI water), then poured onto nutrient agar plates for *φ*X174 and LB agar plates for T4 and incubated overnight at 37°C. Plaques were counted the next day, and the phage

**Fig. 1.**
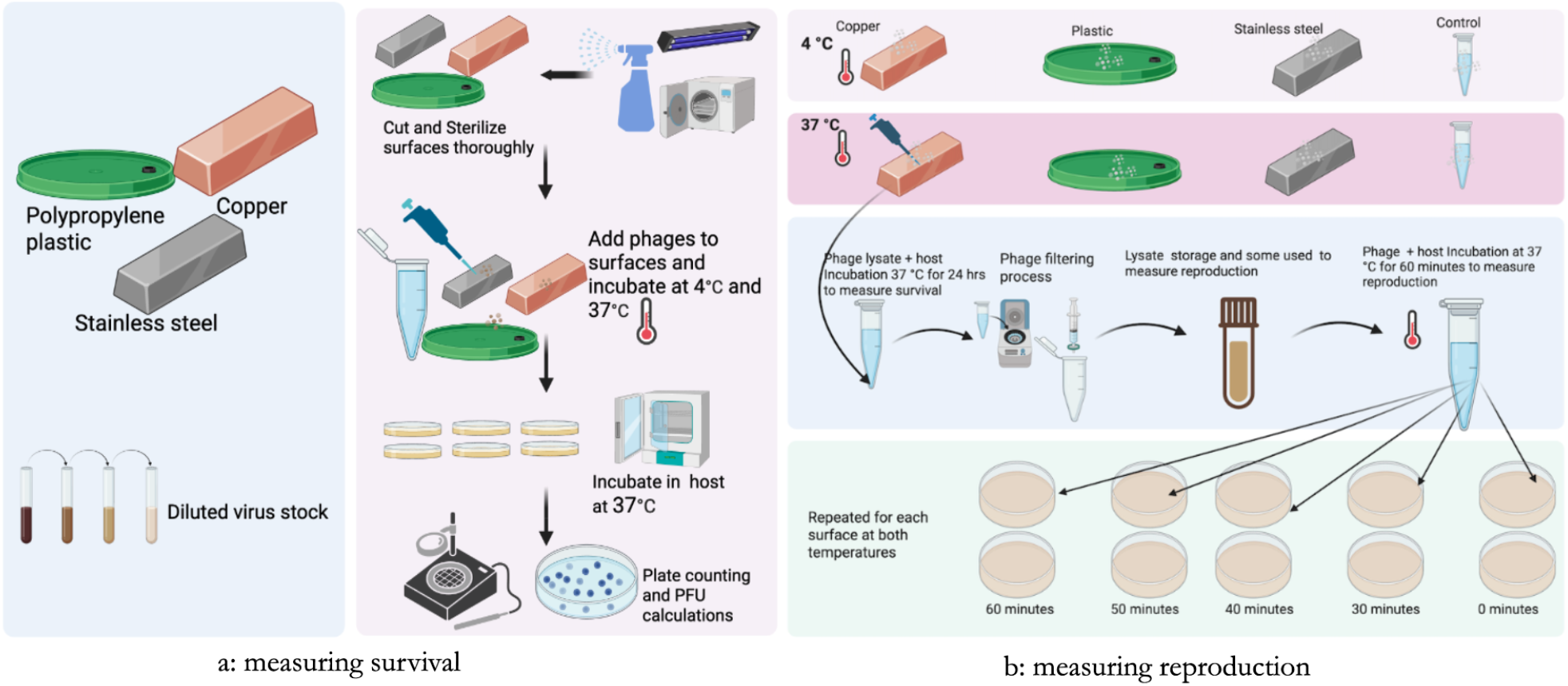
Illustration of experimental workflow. Viruses were quantified for survival and subsequently harvested to measure reproductive efficiency at an hour time point. (a) Surfaces were prepared by thoroughly cleaning, disinfecting, and sterilizing. Viral particles were then suspended on the surfaces and incubated at the appropriate temperature for eight hours. During incubation, viral samples were harvested at 2-hour intervals and then incubated in host cells to measure survival. (b) The phage lysate at the end of eight hours was harvested to measure reproduction. titers were calculated.

### 2.2 Statistical analyses

We applied several statistical methods to investigate differences in T4 and *φ*X174 survival and reproduction across different surfaces and temperatures. We applied non-parametric tests (e.g., Kruskal-Wallis) (*68*) since some groups exhibited constant or near-constant values after 8 hours of exposure on a surface, for example, phages lost viability completely on copper after 8 hours of exposure. To quantify how rapidly loss of viability happened, we fit simple linear regression models of the form:

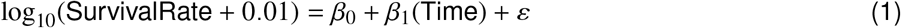

where the slope *ε*_1_ describes the average rate of decline in log-transformed survival. From this, we derive half-lives and percentage decline rates per hour. To understand how phage type and temperature jointly affect survival at the final time point at different surfaces, we applied two-way ANOVA for each surface:

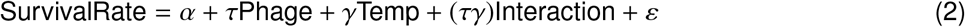

To understand how the two phages reproduced after exposure to these surfaces for eight hours, we applied standard descriptive and comparative statistics. Further, we explored temperature sensitivity on reproduction efficiency. Finally, we used Spearman correlations to understand if there was a relationship between survival and reproduction.

## 3 Results

To investigate how temperature and surface type influence viral stability, we employed two well-characterized bacteriophages, T4 and *φ*X174. Each was exposed to copper, stainless steel, and polypropylene plastic at 4°C and 37°C. Temperature exerted a pronounced effect on viral survival for both phages. Viruses incubated at 37°C experienced greater reductions in viability than those exposed to 4°C on the same surfaces (**Figure** 2, **Figure** S1,**Table** S1). Notably, we observed that viral populations could recover reproductive capacity even after becoming undetectable in survival assays (**Figure** 3, **Figure** S2, **Figure** S3). The relationship between survival and reproduction varied across environments. For example, *φ*X174 exhibited a strong positive correlation between survival and reproduction on copper surfaces, but a negative association between these same traits on polypropylene plastic (**Figure** 4).

**Fig. 2.**
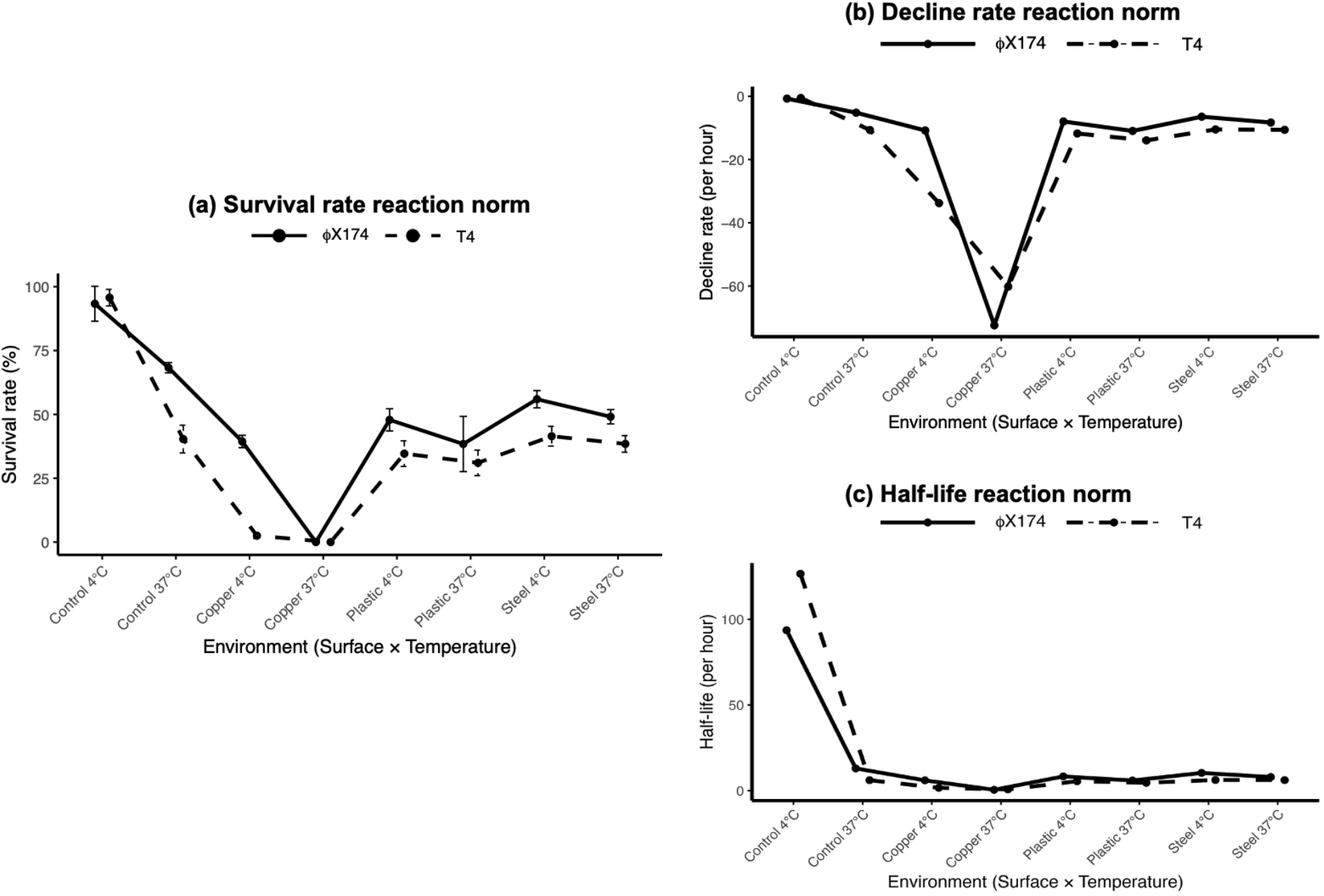
Reaction norms demonstrating how *φ*X174 and T4 viruses survive at three different surfaces: copper, stainless steel, and plastic, exposed to two different temperatures (4°C and 37°C). (a) Survival rates reaction norms. (b) decline rate, and (c) half-life. The survival rate was calculated as the percentage of final virus titers compared to the initial titers. For more details, see **Table** 1. Linear models were used to fit log-transformed survival rates over time to estimate decline rates and half-lives, quantifying how rapidly the two viruses were losing viability at these surfaces and temperatures. Figure (a) shows the percentage decline rates per hour, while Figure (b) shows the half-lives in hours for both phages. Both phages show very high decline rates and short half-lives on copper exposed to higher temperatures. See **Table** 2 for more details.

**Fig. 3.**
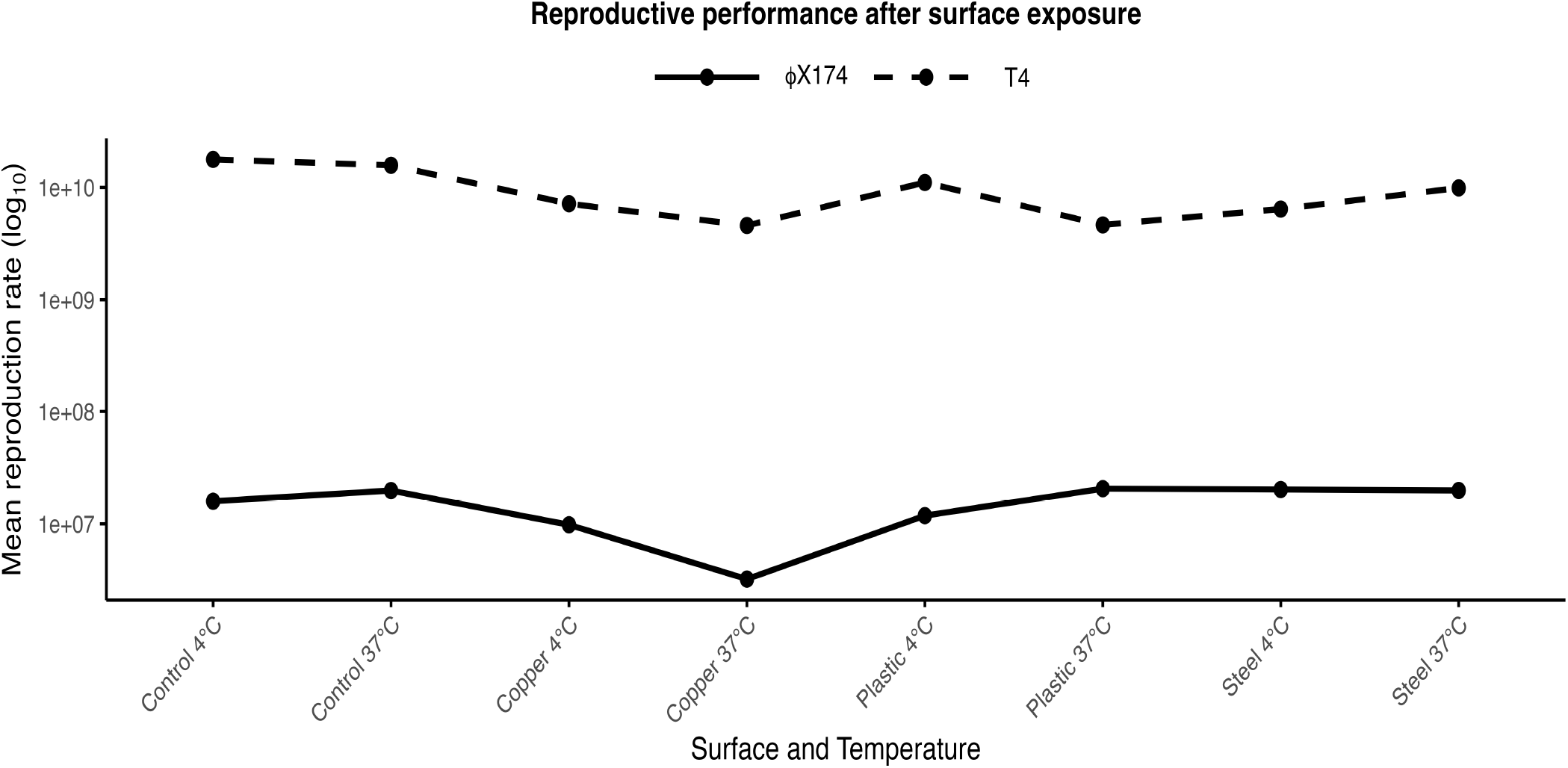
Reaction norms for reproductive performance of *φ*X174 and T4 viruses after exposure to copper, stainless steel, and plastic at two different temperatures: Reproduction was measured by taking viral lysates from the final point of survival experiments and allowing them to grow in fresh media over a period of 60 minutes. See **Figure** 1 for more experimental details. Statistical calculations were done on the reproduction data, and the means from each surface temperature were plotted. See **Table** 4 for additional summary metrics related to reproduction, and **Figure** S2, and **Figure** S3. Reproductive performance decreases for both viruses on copper at high temperature. The effect of temperature on reproduction for both phages varies across surface and temperature. *φ*X174 reproduces at higher temperatures on plastic and stainless steel. T4, however, reproduces relatively poorly at higher temperatures on plastic and relatively well on stainless steel. For more details, see **Table** S2.

**Fig. 4.**
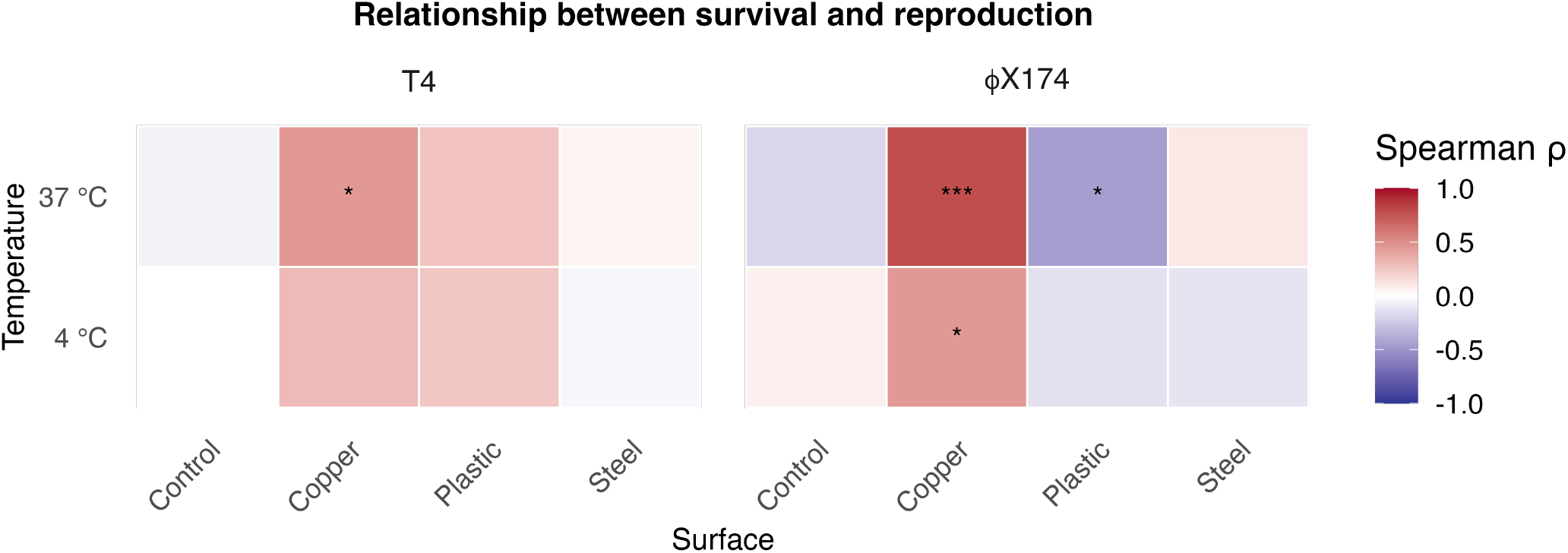
Test of correlation between survival and reproduction across surfaces and temperatures. The relationship between survival (mean surface survival virus titers) and reproduction (virus titers measured during reproduction) for each virus was tested using Spearman’s rank Correlation. A significant positive Spearman’s correlation suggests that as survival performance improves, reproduction also improves, and as it declines, reproduction declines accordingly. Negative correlations imply that conditions that favor survival on one surface and temperature might not favor reproduction; similarly, conditions that favor reproduction on one surface do not warrant survival. On copper at 37°C, *φ*X174 shows a significant correlation between survival and reproduction, suggesting that conditions that favor higher survival in this setting also favor higher reproduction. A positive correlation between survival and reproduction is also observed on copper at 37°C for T4. However, at the same temperature on plastic, this relationship is negative, suggesting that conditions favoring survival here are not suitable for reproduction. For more details, see **Table** 5.

### 3.1 Effects of surfaces and temperature on survival and reproduction

Our results demonstrate that surface type can markedly influence the rate at which phages lose stability, particularly on copper surfaces (**Figure** 2(a), **Table** 1, **Figure** S1). In contrast, both stainless steel and polypropylene plastic supported prolonged viral stability, with minimal temperature effects at 4°C. A Kruskal–Wallis test revealed a significant temperature effect for phages exposed to copper (*p* = 0.0021). For viruses on stainless steel, the identity of phage, rather than temperature, was the significant factor (*p* = 0.0039). On plastic surfaces, neither phage type nor temperature produced a significant effect. Control experiments confirmed that temperature alone influenced stability (*p* = 0.0039), with phages at 4°C exhibiting higher survival than those at 37°C.

**Table 1.** Summary of virus survival rates (% of virus titers at each final point compared to the initial virus titer).

| Surface | PhageType | Temperature(°C) | Initial_Titer | Final_Titer | Survival_Rate (%) | SD |
| --- | --- | --- | --- | --- | --- | --- |
| Control | φX174 | 4 | 1.74e+08 | 1.62e+08 | 93.33 | 6.84 |
| Control | T4 | 4 | 1.46e+10 | 1.40e+10 | 95.72 | 3.23 |
| Control | φX174 | 37 | 1.80e+08 | 1.23e+08 | 68.29 | 2.01 |
| Control | T4 | 37 | 1.56e+10 | 6.27e+09 | 40.34 | 5.44 |
| Copper | φX174 | 4 | 1.48e+08 | 5.82e+07 | 39.39 | 2.42 |
| Copper | T4 | 4 | 1.28e+10 | 3.17e+08 | 2.50 | 1.07 |
| Copper | φX174 | 37 | 1.45e+08 | 0.00e+00 | 0.00 | 0.00 |
| Copper | T4 | 37 | 1.28e+10 | 0.00e+00 | 0.00 | 0.00 |
| Plastic | φX174 | 4 | 1.43e+08 | 6.83e+07 | 47.86 | 4.35 |
| Plastic | T4 | 4 | 1.30e+10 | 4.48e+09 | 34.64 | 5.03 |
| Plastic | φX174 | 37 | 1.49e+08 | 5.75e+07 | 38.40 | 10.81 |
| Plastic | T4 | 37 | 1.28e+10 | 3.95e+09 | 31.04 | 4.99 |
| Steel | φX174 | 4 | 1.53e+08 | 8.53e+07 | 55.95 | 3.39 |
| Steel | T4 | 4 | 1.28e+10 | 5.28e+09 | 41.47 | 3.89 |
| Steel | φX174 | 37 | 1.49e+08 | 7.32e+07 | 49.07 | 2.80 |
| Steel | T4 | 37 | 1.28e+10 | 4.90e+09 | 38.44 | 3.26 |

### 3.2 Survival on surfaces

We used linear models to estimate the rate of loss of viability for T4 and *φ*X174 across surfaces. On stainless steel and plastic, *φ*X174 displayed slightly slower decline rates than T4 at equivalent temperatures **Figure** 2(b), and **Table** 2, suggesting differences in environmental persistence between the two viruses. Both phages exposed to copper at 37°C exhibited pronounced reductions in half-life (**Figure** 2(c), and **Table** 2). *φ*X174 had a half-life of approximately 0.54 hours, while T4 showed near-complete inactivation, indicating a strong thermal effect on copper surfaces. Here we see a very clear example of a crossing reaction norm, where *φ*X174 has a lower decline rate across most surface-temperature combinations. However, on copper, its decline rate is higher than that of T4. In the parlance of reaction norms in quantitative genetics (*69,70*), crossing reaction norms is often indicative of a gene-by-environment interaction, where the rank performance of genotypes is specific to environmental context (*71*). In this study, it indicates that the answer to the question of which virus type (*φ*X174 or T4) is the superior survivor depends on context. *φ*X174 seems specifically susceptible to decay when exposed to copper at 37°C.

**Table 2.** Summary results of the rate of decline analysis by surface, temperature (Temp), and phage type estimated by fitting linear models to log-transformed survival rates over time to estimate slopes, half-lives, and decline rates. Negative slopes indicate a decline in survival rates.

| Surface | Phage | Temp (°C) | mean_slope | sd_slope | mean_r2 | n | half_life | decline rate(hr) |
| --- | --- | --- | --- | --- | --- | --- | --- | --- |
| Control | φX174 | 4 | -0.0032 | 0.0026 | 0.348 | 3 | 93.63 | -0.74 |
| Control | T4 | 4 | -0.0024 | 0.0041 | 0.189 | 3 | 126.59 | -0.55 |
| Control | φX174 | 37 | -0.0231 | 0.0023 | 0.940 | 3 | 13.01 | -5.19 |
| Control | T4 | 37 | -0.0490 | 0.0072 | 0.987 | 3 | 6.14 | -10.67 |
| Copper | φX174 | 4 | -0.0497 | 0.0042 | 0.961 | 3 | 6.06 | -10.82 |
| Copper | T4 | 4 | -0.1791 | 0.0191 | 0.782 | 3 | 1.68 | -33.79 |
| Copper | φX174 | 37 | -0.5594 | 0.0029 | 0.791 | 3 | 0.54 | -72.42 |
| Copper | T4 | 37 | -0.4000 | 0.0000 | 0.500 | 3 | 0.75 | -60.19 |
| Plastic | φX174 | 4 | -0.0361 | 0.0044 | 0.877 | 3 | 8.34 | -7.98 |
| Plastic | T4 | 4 | -0.0544 | 0.0062 | 0.899 | 3 | 5.53 | -11.77 |
| Plastic | φX174 | 37 | -0.0504 | 0.0132 | 0.958 | 3 | 5.97 | -10.97 |
| Plastic | T4 | 37 | -0.0652 | 0.0104 | 0.981 | 3 | 4.62 | -13.94 |
| Steel | φX174 | 4 | -0.0290 | 0.0030 | 0.960 | 3 | 10.38 | -6.46 |
| Steel | T4 | 4 | -0.0481 | 0.0046 | 0.931 | 3 | 6.25 | -10.49 |
| Steel | φX174 | 37 | -0.0376 | 0.0038 | 0.974 | 3 | 8.00 | -8.30 |
| Steel | T4 | 37 | -0.0487 | 0.0039 | 0.974 | 3 | 6.18 | -10.61 |

### 3.3 Combined effects of phage type and temperatures on survival across surfaces

We performed a two-way ANOVA to assess the combined effects of phage type and temperature on viral survival across surfaces. On copper, both factors had significant effects, and their interaction was highly significant, indicating a strong synergistic influence on phage stability (**Table** 3). On stainless steel, phage type was significant and temperature exhibited a moderate effect, though their interaction was not significant. No significant effects were detected on plastic surfaces. In the control experiment, both temperature and phage type showed a highly significant interaction (**Table** 3).

**Table 3.** Two-way ANOVA results by surface: Comparing interaction effects.

| Surface | term | statistic | p.value | significance |
| --- | --- | --- | --- | --- |
| Control | Phage Type | 21.59 | 0.0017 | ** |
| Control | Phage × Temperature | 30.41 | 0.0006 | *** |
| Control | Temperature | 213.68 | 0.0000 | *** |
| Copper | Phage Type | 582.12 | 0.0000 | *** |
| Copper | Phage × Temperature | 582.12 | 0.0000 | *** |
| Copper | Temperature | 750.71 | 0.0000 | *** |
| Plastic | Phage Type | 6.84 | 0.0309 | * |
| Plastic | Phage × Temperature | 0.55 | 0.4777 | ns |
| Plastic | Temperature | 2.75 | 0.1356 | ns |
| Steel | Phage Type | 42.00 | 0.0002 | *** |
| Steel | Phage × Temperature | 0.99 | 0.3491 | ns |
| Steel | Temperature | 6.52 | 0.0340 | * |

### 3.4 Reproductive performance following surface exposure

After examining the free-living survival of the phages, we measured how survivors reproduced following exposure to different surface and temperature conditions. Both T4 and *φ*X174 replicated within one hour and exhibited substantial increases across surfaces (**Figure** 3, **Figure** S2). The normalized curves for *φ*X174 showed robust increases, exceeding 100-fold within an hour on plastic surfaces and in the control (**Figure** S3). In contrast, T4 displayed more modest increases, particularly at 37°C, where its titers rose only slightly relative to initial values (**Figure** S3). Shifting temperatures from 4°C to 37°C also affected viral reproduction, although this effect was different across surfaces (**Figure** 3). Additionally, we calculated final virus titer ratios, growth-phase ratios, and temperature effects within one hour of reproduction to quantify the observed patterns between the two phages (**Table** 4). Compared to *φ*X174, T4 generally reached high titers. However, on copper at 37°C, all phages completely lost viability, consistent with the strong inactivating effect of this environment.

**Table 4.**
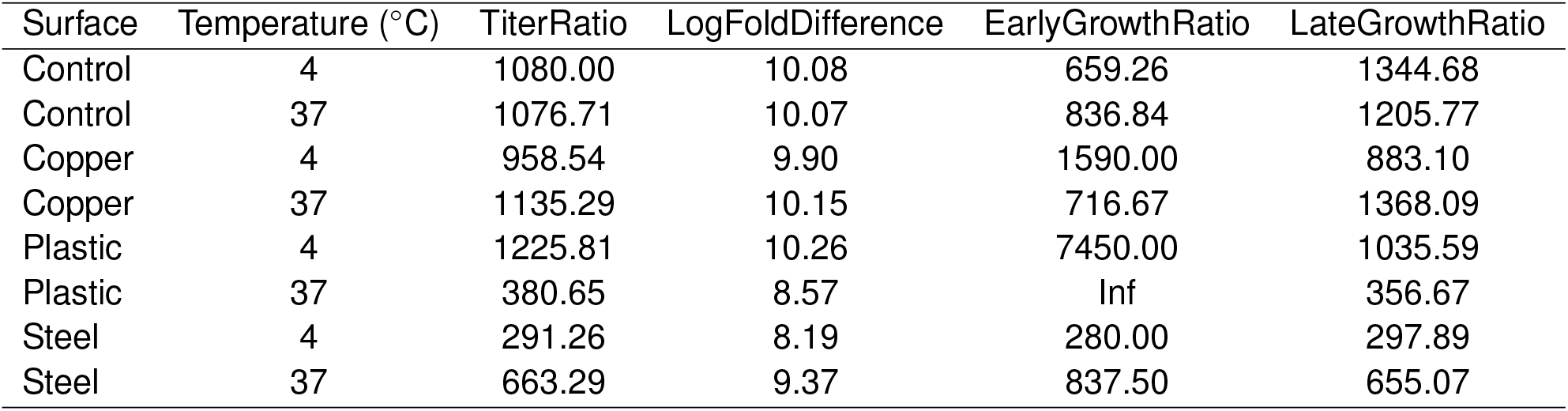
Titer Ratios (T4/*φ*X174) reflecting reproductive efficiency after exposure to different surfaces and temperatures. We split reproduction into two phases: EarlyGrowth (0-30 minutes), and LateGrowth (30-60 minutes), and compared the growth ratios. Early growth ratios for T4 were high. For example, T4 on plastic reached extremely high numbers at 37°C as indicated by “Inf” and 1225 at 4°C. See **Table** S1 for additional details.

### 3.5 Reproductive rates and temperature sensitivity

Temperature effects on phage reproduction were not uniform across phages. When surface temperature increased from 4°C to 37°C, *φ*X174 showed modest declines in reproductive efficiency, with ratios near or below 1, and the largest reduction was observed on stainless steel (approximately 0.77). In contrast, T4 exhibited greater variability across surfaces, including a marked increase in reproductive efficiency on stainless steel (approximately 1.75) ((**Table** S1,and **Table** S2).

### 3.6 Relationship between survival and reproduction

We next examined how phage survival related to reproductive performance across surfaces using Spearman’s rank correlation. The relationship between survival and reproduction was not consistent across contexts: higher survival did not necessarily predict faster reproduction, and vice versa. For *φ*X174 at 37°C on copper, we observed a strong positive correlation between survival and reproduction (*ρ* = 0.784, *p* = 3.5 10^−6^), suggesting that under these conditions, stability and reproduction were linked (**Figure** 4, **Table** 5). However, for *φ*X174 on plastic at the same temperature, the correlation was negative (*ρ* = 0.451, *p* = 0.235), indicating that conditions favoring survival may not favor reproduction. T4 at 37°C also displayed a positive association between survival and reproduction (*ρ*= 0.451, *p* = 0.0235). In other contexts, relationships were weak or non-significant, suggesting that the connection between phage persistence and reproductive potential is highly context-dependent.

**Table 5.**
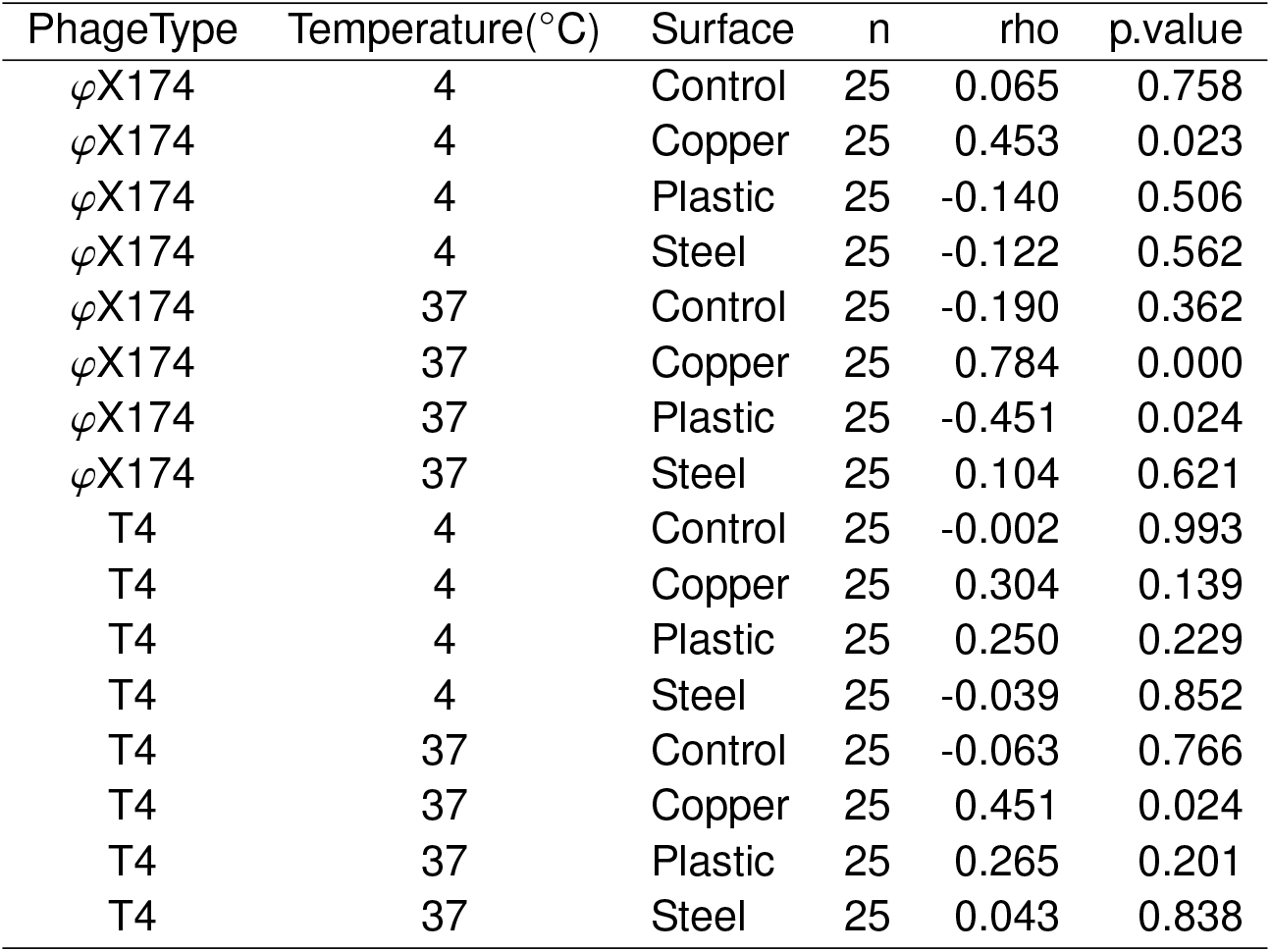
Spearman correlation results between survival and reproduction.

| PhageType | Temperature(°C) | Surface | n | rho | p.value |
| --- | --- | --- | --- | --- | --- |
| φX174 | 4 | Control | 25 | 0.065 | 0.758 |
| φX174 | 4 | Copper | 25 | 0.453 | 0.023 |
| φX174 | 4 | Plastic | 25 | -0.140 | 0.506 |
| φX174 | 4 | Steel | 25 | -0.122 | 0.562 |
| φX174 | 37 | Control | 25 | -0.190 | 0.362 |
| φX174 | 37 | Copper | 25 | 0.784 | 0.000 |
| φX174 | 37 | Plastic | 25 | -0.451 | 0.024 |
| φX174 | 37 | Steel | 25 | 0.104 | 0.621 |
| T4 | 4 | Control | 25 | -0.002 | 0.993 |
| T4 | 4 | Copper | 25 | 0.304 | 0.139 |
| T4 | 4 | Plastic | 25 | 0.250 | 0.229 |
| T4 | 4 | Steel | 25 | -0.039 | 0.852 |
| T4 | 37 | Control | 25 | -0.063 | 0.766 |
| T4 | 37 | Copper | 25 | 0.451 | 0.024 |
| T4 | 37 | Plastic | 25 | 0.265 | 0.201 |
| T4 | 37 | Steel | 25 | 0.043 | 0.838 |

## 4 Discussion

To investigate how abiotic stressors—surfaces and temperature—shape viral persistence and reproduction, we challenged bacteriophages T4 and *φ*X174 across copper, stainless steel, and polypropylene at 4°C and 37°C. Viral survival was strongly influenced by both temperature and surface composition, and these effects were frequently non-additive. Although the effects of these stressors were more straightforward with respect to reproduction, the relationship between stability and reproductive efficiency did not reduce to simple patterns but instead reflected interactions among surface composition, thermal context, and virus type.

When comparing T4 with *φ*X174, one could ask the simple question: “Which has higher free-living survival?” Our results suggest that it depends. In our results, *φ*X174 has a higher survival across most temperatures and surface compositions. But the complete answer is context dependent: *φ*X174 has a higher free-living survival in most surface-temperature combinations, as dictated by several metrics (decline rate, half-life). But at 37°C on copper, it actually has a lower survival rate than T4. The crossing reaction norms of the survival curves across surface-temperature combinations is indicative of what we label a “virus type by environment interaction,” which we consider a kind of genotype by environment interaction (G× E). And our study highlights that reproduction is much more resistant to surface-temperature combinations, with T4 having a higher mean reproduction rate in all environments tested. This can also be framed in terms of environmental robustness with respect to reproduction, a measure of the invariance in the performance of a particular trait or organism (*72,73*). We can say that both viruses demonstrate environmental robustness with respect to survival, with T4 having a higher average reproduction.

Looking across surface compositions and temperatures, several consistent patterns emerge: every virus type has its lowest survival on copper surfaces. In particular, warmer temperatures accelerated viral decay relative to cooler conditions. These results extend previous demonstrations of copper’s antimicrobial activity to varying temperatures, and align with direct evidence that copper and copper alloys rapidly inactivate viruses on contact, whereas stainless steel often facilitates longer persistence (*36,74–76*). Similarly, these results are in line with a long literature on the role of thermal stressors on viral performance.

### Implications for infectious disease ecology

Beyond free-living survival, our experiments show that post-exposure reproduction is not a simple function of residual viability. The correlations between survival and reproduction varied in sign and magnitude between environments, implying that stability and reproductive output represent partially decoupled axes of fitness shaped by the environmental context. This is a significant outcome when we consider the larger literature on infectious disease ecology and life history theory that has long examined the constraints governing relationships between traits such as virulence and transmission, survival, and reproduction (*51,55,58,59,77,78*). Our findings demonstrate that survival and reproduction are largely decoupled and that their relationship is specific to certain surface-temperature combinations. This supports modern notions that trait relationships in pathogen-host systems are idiosyncratic and context-dependent.

### Implications for biomedicine and epidemiology

Although our study uses model bacteriophages, the findings have implications for fomite-mediated transmission and for applications such as phage therapy for medical devices and implants. Questions about how pathogens survive on materials underlie efforts to reduce surface contamination through the use of antimicrobial copper materials (*79*).

The findings also resonate with evolutionary responses to metal stress in microbes, as was demonstrated in a recent study on long-term copper exposure in *Escherichia coli* (*80*). This invites future work asking whether repeated exposure to surfaces can drive viral adaptations in persistence or replication, and whether such adaptations reshape the coupling between survival and reproduction. With regard to epidemiology, our findings add further nuance to our conception of the forces shaping fomite-mediated transmission. Models of fomite-mediated transmission often include a decay rate parameter to describe how pathogen populations decline (*81,82*). Our findings suggest that models (conceptual, computational, etc.) must consider how abiotic environments interact in shaping this decay parameter, as they can facilitate more accurate analysis and interventions. Furthermore, public health strategies that aim to intervene in environmental transmission, such as the use of copper surfaces in clinical settings, must consider how these effects are mediated by temperature.

### Limitations

Our study has several limitations. We used bacteriophages as model systems. Although they offer valuable mechanistic insights, they differ from human and animal viruses in important ways. In addition, the time frame of our experiments was restricted to eight hours, which may not capture longer-term persistence dynamics. Lastly, we measured survival and reproduction sequentially rather than simultaneously, which may obscure finer-scale temporal dynamics. Future work should extend these experiments to longer durations, incorporate a broader range of temperatures and humidity conditions, and test additional viral systems, particularly enveloped and RNA viruses of public health importance.

### Conclusion

In conclusion, our findings demonstrate that viral survival and reproduction on abiotic surfaces are shaped by complex, context-dependent interactions between environmental factors. By disentangling how temperature and surface type jointly influence these traits, we provide a framework for better understanding fomite-mediated transmission and for integrating environmental data into models of pathogen ecology and evolution.

## Acknowledgments

The authors thank V. Ezenwa, N. Grubaugh, P. Pennings, P. Turner, members of the OgPlexus for helpful discussions on the topic, and the Yale University Statistics clinic. The authors thank the Yale Quantitative Biology Institute for an invitation to a seminar in which the ideas in this manuscript were discussed.

## Author contributions

KK and CBO conceived the study; KK and TJ performed the experiments; KK, AS, and CBO analyzed and interpreted data; KK and CBO wrote the original and revised versions of the manuscript.

## Data and materials availability

Data and code can be found on Github: https://github.com/OgPlexus/Surface1.

