## Supplemental material for "Surface-by-temperature interactions shape the relationship between viral free-living survival and reproduction"

### Supplementary Material

#### Notes on supplementary material

Here we provide a series of figures and tables that support findings from the main text.

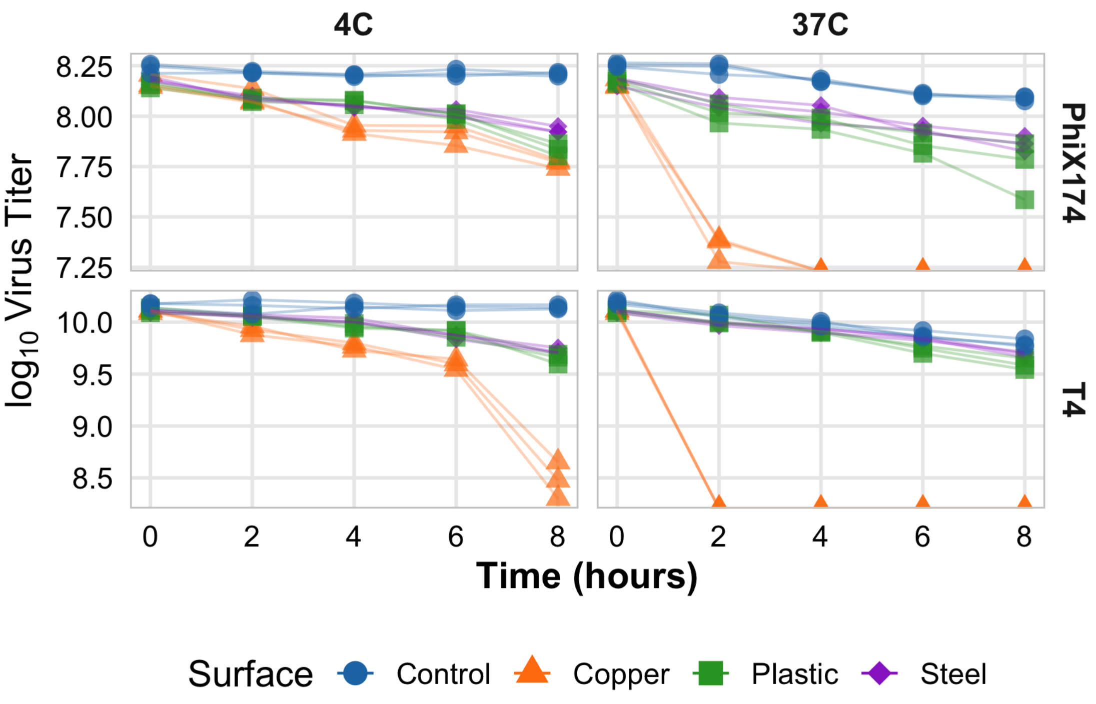


**Fig. S1.** Changes in survival rates for T4 and $\varphi$X174 over time on different surfaces. Each panel represents a distinct surface type, with lines corresponding to mean survival trajectories at 4°C and 37°C. The y-axis is shown on a logarithmic scale to highlight differences at low survival levels. Both phages exhibit distinct survival dynamics across surfaces and temperatures, with $\varphi$X174 and T4 displaying characteristic patterns of stability under each condition.

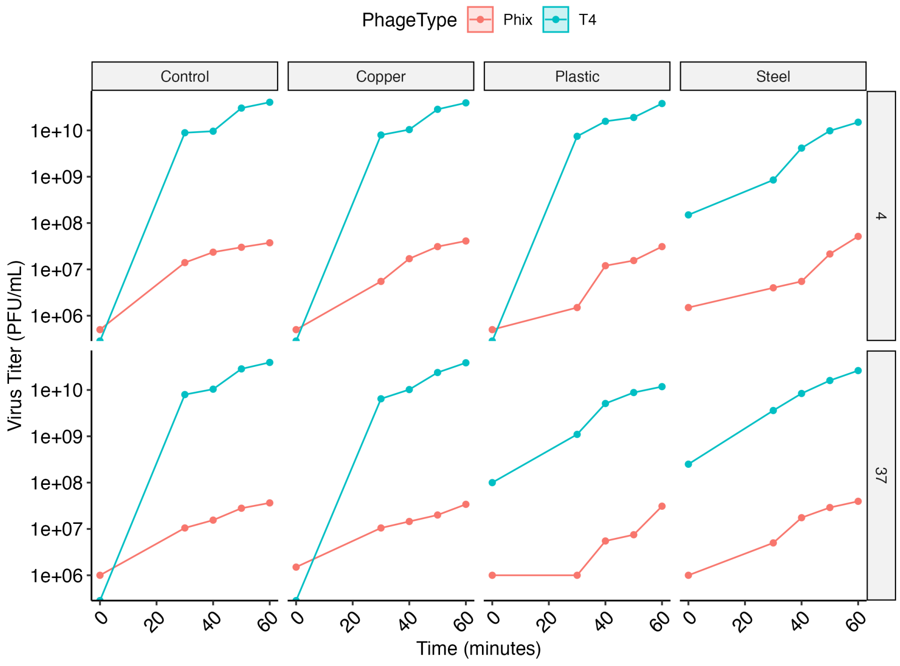


**Fig. S2.** Mean virus titers plotted over time (minutes). Both T4 and $\varphi$X174 replicate substantially within 60 minutes. However, T4 virus titers are higher than $\varphi$X174 across all surfaces for both 4°C and 37°C.

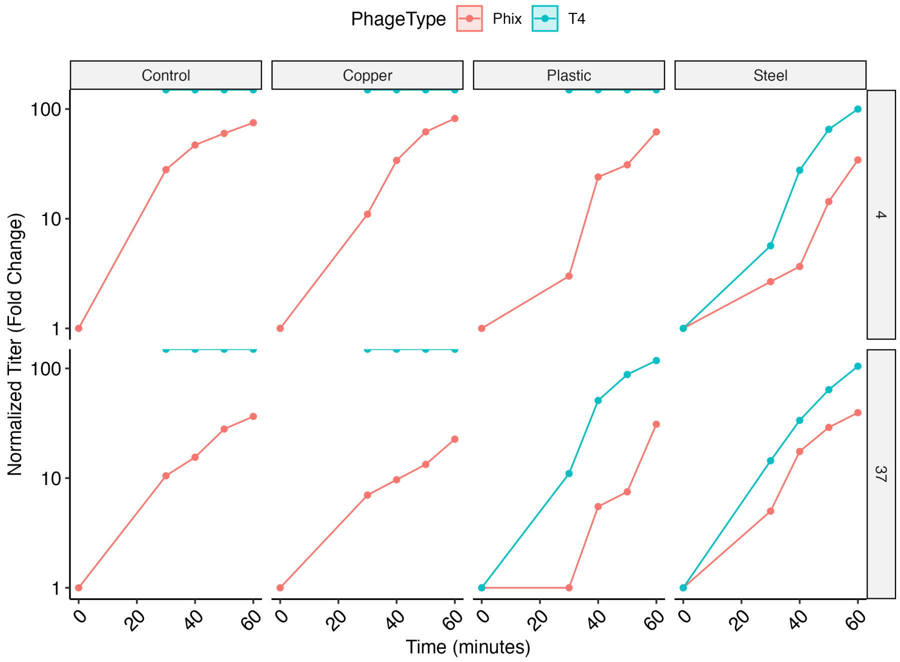


**Fig. S3.** In these normalized curves, each phage's growth is expressed relative to its own initial titer (fold-change): $\varphi$X174 shows a robust fold-increase, particularly at 4°C on plastic and control surfaces.

**Table S1.** Virus titers for reproduction are divided into Early (0-30 minutes) and Late ( 30-60 minutes) Growth phases.

| Surface | Temperature(°C) | EarlyGrowth$\varphi$X174 | EarlyGrowthT4 | LateGrowth$\varphi$X174 | LateGrowthT4 | EarlyGrowthRatio | LateGrowthRatio |
|---|---|---|---|---|---|---|---|
| Control | 4 | 4.50e+05 | 2.97e+08 | 7.83e+05 | 1.05e+09 | 659.26 | 1344.68 |
| Control | 37 | 3.17e+05 | 2.65e+08 | 8.67e+05 | 1.04e+09 | 836.84 | 1205.77 |
| Copper | 4 | 1.67e+05 | 2.65e+08 | 1.18e+06 | 1.04e+09 | 1590.00 | 883.10 |
| Copper | 37 | 3.00e+05 | 2.15e+08 | 7.83e+05 | 1.07e+09 | 716.67 | 1368.09 |
| Plastic | 4 | 3.33e+04 | 2.48e+08 | 9.83e+05 | 1.02e+09 | 7450.00 | 1035.59 |
| Plastic | 37 | 0.00e+00 | 3.33e+07 | 1.00e+06 | 3.57e+08 | Inf | 356.67 |
| Steel | 4 | 8.33e+04 | 2.33e+07 | 1.58e+06 | 4.72e+08 | 280.00 | 297.89 |
| Steel | 37 | 1.33e+05 | 1.12e+08 | 1.15e+06 | 7.53e+08 | 837.50 | 655.07 |

**Table S2.** Temperature sensitivity analysis. Temperature effects on virus reproduction vary depending on virus type and surface.

| PhageType | Surface | TempEffect | LogTempEffect |
|---|---|---|---|
| $\varphi$X174 | Control | 0.97 | -0.04 |
| $\varphi$X174 | Copper | 0.83 | -0.27 |
| $\varphi$X174 | Plastic | 1.00 | 0.00 |
| $\varphi$X174 | Steel | 0.77 | -0.38 |
| T4 | Control | 0.97 | -0.04 |
| T4 | Copper | 0.98 | -0.03 |
| T4 | Plastic | 0.31 | -1.69 |
| T4 | Steel | 1.75 | 0.80 |
